# Immune-induced TCR-like antibodies regulate specific T cell response

**DOI:** 10.1101/2025.02.03.636362

**Authors:** Kazuki Kishida, Keisuke Kawakami, Hiroaki Tanabe, Wataru Nakai, Koji Yonekura, Shigeyuki Yokoyama, Hisashi Arase

## Abstract

Antigen-specific regulation of T cell response is crucial for limiting hyperimmune response. In this study, we discovered that antibodies specific to the antigen peptide-MHC class II complex are produced during helper T cell responses to various antigens. These antibodies specifically inhibited T cell receptor (TCR) recognition of MHC class II molecules presenting specific antigen peptide. We have termed these antibodies ‘immune-induced TCR-like antibodies’ (iTabs). Immunization with peptides containing flanking residues induced iTabs, whereas immunization with peptides lacking flanking residues did not. Furthermore, immunization of iTab-inducible peptide as well as iTab treatment suppressed autoimmune disease development. Our findings provide a strategy for suppressing antigen-specific helper T cell responses using specific peptides, potentially controlling autoimmune diseases.

## Main text

Immunization with antigens triggers antigen-specific T cell and B cell activation, leading to antibody production against the antigens. Major histocompatibility complex class II molecules (MHC-II) are crucial for adaptive immunity, presenting peptide antigens to helper T cells. Certain MHC-II alleles are strongly associated with the risk of autoimmune diseases and allergies (*1, 2*). Thus, controlling T cell receptor (TCR) recognition of peptides presented on MHC-II is vital for controlling the immune response. Antibodies (Abs) that recognize the peptide-MHC-II complex, exhibiting similar specificity to TCRs, can be generated by immunizing with the peptide-MHC-II complex, providing a tool for analyzing antigen presentation (*3, 4*) and blocking TCR recognition (*5, 6*). However, it has been generally thought that these TCR-like Abs are not typically produced during normal immune responses.

The peptide-binding groove of MHC-II is open at both ends, resulting in fewer restrictions on the peptide length (*7*). Indeed, not only peptides but also whole proteins can be presented on MHC-II (*8–11*). Although minimal peptides suffice for helper T cell recognition and are commonly used in studies, naturally presented peptides on MHC-II often possess long flanking residues (FR), whose roles in immunity are not fully understood (*12–14*). This study reveals that immune responses to antigens induce significant amounts of TCR-like Abs that suppress antigen-specific helper T cell responses by blocking TCR recognition of the peptide-MHC-II complex. Furthermore, we demonstrate that peptides with FR but not minimal peptides, also induce TCR-like Abs. These findings provide a novel strategy for regulating specific immune responses.

### Immunization with protein antigen induces Abs against the peptide-MHC-II complex

We examined whether a general antigen-specific immune response produces Abs against the peptide-MHC-II complex. B10.A mice (I-A^k^) were immunized with hen egg lysozyme (HEL) protein, and serum Ab binding to 30-amino-acid overlapping HEL peptides pulsed onto 293T cells expressing MHC-II, I-A^k^, was analyzed. Cells pulsed with HEL_41-70_ and HEL_51-80_ peptides, both containing the major T cell epitope, HEL_48-61_ (*15*), were recognized by the serum Abs (**Fig. 1A**). In contrast, cells pulsed with HEL_48-61_, containing only the major T cell epitope, were not recognized by the Abs, whereas cells pulsed with the HEL_48-61_ were recognized by the monoclonal Ab, Aw3.18, specific for HEL peptides presented on MHC-II (*3*), indicating that these HEL peptides are equally presented on MHC-II (**Fig. 1B**). Similar to the Aw3.18 Ab, a single-chain TCR-Fc fusion protein of the 3A9 TCR, specific to HEL_48-61_ peptides presented on MHC-II (3A9 TCR-Fc)(*16*), also bound to HEL_48-61_ peptide-pulsed cells in a dose-dependent manner (**fig. S1 and Fig. 1B**). When HEL_48-61_ peptides with additional N-terminal (HEL_41-61_) or C-terminal (HEL_48-70_) FR were pulsed, the serum Abs bound to HEL_48-70_ with a C-terminal, but not N-terminal FR. At least two tryptophan residues at the C-terminal FR are necessary for the Ab binding (**Fig. 1C**). These data indicated that immunization with HEL protein induces the Abs that recognize HEL peptides with C-terminal FR presented on MHC-II expressing cells.

Next, we investigated whether the Abs produced by HEL protein immunization recognize the FR of HEL peptides alone or the peptide-MHC-II complex. HEL protein-immunized serum Abs were absorbed on HEL protein- or HEL_48-64_ peptide-bound Sepharose™ beads (**Fig. 1D**). Antibody titers against the HEL protein or the HEL_48-64_ peptide itself but not the HEL peptides presented on MHC-II were decreased upon absorption (**Fig. 1D**). This indicated that Abs specific to the peptide-MHC-II complex were generated by HEL protein immunization. Because these Abs recognize both peptides and MHC-II similar to TCRs, we termed these Abs as **immune**-induced TCR-like antibodies (iTabs). Interestingly, iTabs titers were decreased from two weeks post-immunization, whereas anti-HEL Ab titers were increased (**Fig. 1E**), suggesting that the dynamics of iTab production differ from those of a general Ab response.

Since both TCRs and iTab bind to peptides presented on MHC-II, we investigated whether iTabs block TCR recognition. 3A9 TCR-Fc binding to HEL_48-64_-loaded cells was blocked dose-dependently by iTabs from HEL protein-immunized mice (**Fig. 1F**). However, the binding of TCR-Fc to HEL_48-61_-loaded cells was not blocked by the iTab. This indicated that iTabs that block anti-HEL T cell recognition were produced during the immune response to HEL protein. Mass spectrometry analysis confirmed that HEL peptides with C-terminal FR such as HEL_48-63_ and HEL_48-64_ are presented on MHC-II (*15*). Immunization of B10 (I-A^b^), B10.D2 (I-A^d^), and B10.S (I-A^s^) congenic mice with HEL protein induced Abs against the HEL protein itself. However, serum Abs did not bind to HEL_48-64_ pulsed cells, thus not blocking 3A9 TCR-Fc binding (**Fig. 1G**). These data indicate that the iTabs exhibit MHC-restricted recognition similar to TCRs.

### Peptide FR is required for iTab production

We next examined whether HEL peptide immunization induces iTabs like HEL protein immunization. Immunization with HEL_48-70_ or HEL_48-64_ (FR^+^ HEL), containing C-terminal FR, but not HEL_40-61_ or HEL_48-61_ (FR^−^ HEL), lacking C-terminal FR, induced iTabs that bind to cells pulsed with HEL_48-64_ or HEL_48-70_ (**Fig. 2A**). iTabs did not bind to cells pulsed with HEL_40-61_ (**Fig. 2A**). These data indicated that C-terminal FR are essential for iTabs induction. To investigate the role of FR in iTab production, we analyzed another peptide antigen from *Schistosoma mansoni,* Sm-P40(*17*). Immunization with Sm-P40_234-246_ induced iTabs against the Sm-P40_234-246_-MHC-II complex, but not the Sm-P40_237-249_-MHC-II complex (**Fig. 2B**). On the other hand, immunization with Sm-P40_237-246_ and Sm-P40_237-249_ did not induce iTabs against the Sm-P40-MHC-II complex. This suggests that N-terminal FR are necessary for inducing iTabs, and C-terminal FR are not required unlike the HEL peptides.

To analyze the role of C-terminal and N-terminal FR in inducing iTabs, we generated recombinant HEL or Sm-P40 peptides containing C-terminal WWA FR or N-terminal PKS FR (**fig. S2A**). HEL peptide with N-terminal PKS FR induced Abs against the Sm-P40_234-246_-MHC- II complex (**Fig. 2B**). Similarly, Sm-P40 peptide with the C-terminal WWA FR elicited Abs against the HEL_48-64_-MHC-II complex (**Fig. 2B**). These results indicated that the FR are required for iTabs production, although the requirement of N- or C-terminal FR depends on the antigen.

Next, we analyzed the role of the FR on the naturally processed peptides on MHC-II. Invariant chain (Ii) and H2-M are essential molecules for peptide presentation by MHC-II (*7*). We generated Ii or H2-Mα knockout LK35.2 cell lines expressing I-A^k^ (**fig. S2B**). When wild-type (WT) LK35.2 cells were incubated with HEL protein, Aw3.18 Ab and 3A9 TCR-Fc bound to the cells, indicating that naturally processed HEL peptides were presented on MHC-II (**fig. S2C**). Moreover, serum iTabs from mice immunized with FR^+^ HEL peptide bound to WT LK35.2 cells pulsed with HEL protein, but serum Abs from mice immunized with FR^−^ HEL peptide did not (**Fig. 3A**). In contrast, when Ii or H2-Mα knockout LK35.2 cells were incubated with HEL protein, Aw3.18, 3A9 TCR-Fc and the serum iTabs did not bind to the cells (**fig. S2C, and Fig. 3A**). These results indicated that peptides with FR are naturally presented on MHC-II during normal antigen processing in APCs and are recognized by iTabs.

### iTabs suppress antigen-specific helper T cell response

iTabs induced by HEL protein immunization blocked 3A9 TCR-Fc binding to HEL peptide presented on MHC-II, suggesting that iTabs suppress antigen-specific helper T cell responses. iTabs induced by immunization with FR^+^ HEL peptide but not FR^−^ HEL peptide also blocked 3A9 TCR-Fc binding to HEL protein pulsed APCs (**fig. S2D**). Furthermore, activation of 3A9 TCR GFP reporter cells was blocked by serum iTabs induced by FR^+^ HEL peptide immunization (**Fig. 3B and 3C**). To further analyze the fine specificity of iTabs, we generated a monoclonal iTab (clone 11-72) from FR^+^ HEL peptide-immunized mice (**fig. S3A**). The 11-72 mAb recognized the HEL peptide-MHC-II complex but not the MHC-II (**fig. S3B**) or the peptide alone (**fig. S3C**) and blocked 3A9 TCR reporter cell activation (**Fig. 3C**). Notably, 11- 72 mAb blocked the binding of 3A9 TCR-Fc stronger than Aw3.18 mAb that recognizes HEL peptides presented on MHC-II (**fig. S3D**). These results suggest that the iTabs are involved in the inhibition of antigen-specific T cell activation.

To analyze the function of iTabs *in vivo*, FR^+^ HEL peptide immunized mice were treated with anti-HEL iTab, 11-72. T cells from anti-HEL iTab-treated mice showed a reduced IL-2 production compared with T cells from control mAb-treated mice (**Fig. 3D**). Similarly, the delayed-type hypersensitivity (DTH) response was also suppressed in anti-HEL iTab-treated mice (**Fig. 3E**). These results suggested that iTabs regulate antigen-specific immune response *in vivo*. To avoid HEL-specific T cell activation by FR^+^ HEL peptide immunization, we generated a FR^+^ HEL peptide in which HEL-specific TCR binding residues were mutated (HEL_Y53A-L56A_) (**fig. S2E**)(*18*) and analyzed iTab production. Surprisingly, the FR^+^ HEL_Y53A- L56A_ peptide also induced iTabs similar to wild-type FR^+^ HEL peptide (**Fig. 3F and fig. S2F**). Furthermore, the iTabs blocked 3A9 TCR-Fc binding (**fig. S2G**). Next, we analyzed the effect of the FR^+^ HEL_Y53A-L56A_ peptide immunization on the response to wild-type HEL peptide. Mice were immunized with FR^+^ or FR^−^ HEL_Y53A-L56A_ peptide, followed by wild-type FR^+^ HEL peptide immunization. CD4^+^ T cells from FR^+^ HEL_Y53A-L56A_ peptide immunized mice showed reduced IL-2 production and proliferation compared with CD4^+^ T cells from FR^−^ HEL_Y53A-L56A_ peptide immunized mice (**Fig. 3G**). These data suggested that HEL-specific iTab suppresses the activation of HEL-specific T cell response *in vivo*.

### The overall structure of the MHC-II–HEL peptide–iTab Fab ternary complex

We determined the cryo-EM structure of the MHC-II-HEL peptide iTab Fab ternary complex at a resolution of 3.09-Å (**Fig. 4A**). The data collection and refinement statistics are provided (**fig. S4** and **Table S1**). The structure of the MHC-II–HEL peptide–iTab Fab ternary complex reveals that the Fab fragment binds to the HEL peptide binding groove of MHC-II. The overall structure of MHC-II exhibits excellent superimposition with its previously reported structure (PDB code 1IAK,(*19*)) (**Fig. 4C**). The root mean square deviation (r.m.s.d.) value for the main chain Cα atoms between the MHC-II-HEL peptide-iTab Fab ternary complex and the MHC-II- HEL peptide complex (PDB code 1IAK) is 1.076 Å.

### Binding interfaces for iTab and the MHC-II-HEL peptide complex

Remarkably, the C-terminal FR of the HEL peptide is recognized by the Fab fragment, and its conformation within the MHC-II-HEL peptide-iTab Fab ternary complex differs from that observed in the previously reported structure of the MHC-II-HEL peptide binary complex (PDB code 1IAK,(*19*)) (**Fig. 4B**). Our study elucidates the Fab binding modes of the C-terminal FR of the HEL peptide and MHC-II within the ternary complex.

Firstly, we investigated the 11-72 mAb binding modes of W62 and W63, the C-terminal FR of the HEL peptide, because peptide mutation analysis has demonstrated the essentiality of W62 and W63 for antibody binding (**fig. S5A**). Within the complex, W62 of the HEL peptide is spatially proximate to Y91, S92, R93, Y94, and P95 of the light chain and W48, N59, and Y106 of the heavy chain (**Fig. 4C**). Similarly, W63 of the HEL peptide is in close proximity to Y91 of the light chain (L) and Y34, S102, S103, Y104, Y105, and Y106 of the heavy chain (H). The indole ring of W63 forms a hydrogen bond with the main-chain carbonyl group of S102/H (**Fig. 4D**).

The MHC-II protein’s α chain (I-A^k^α) is recognized by the light chain of the Fab fragment (**Fig. 4E**). Specifically, T69 of I-A^k^α (T69/α) forms hydrogen bonds with Q27/L, while H72/α engages in hydrophobic interactions with R93/L (**Fig. 4E**). Conversely, the β chain (I-A^k^β) interacts with both the light and heavy chains (**Fig. 4F**). The main chain of E59/β and the side chain of Y60/β participate in hydrophobic interactions with Y104/H. The side chain of K63/β forms hydrophobic interactions with W50/L and Y104/H. The side-chain amino group of Q64/β forms hydrogen bonds with the main-chain carbonyl group of Y104/H. Finally, the side chain of Y65/β establishes hydrophobic contacts with the main chain of V29/L and G30/L (**Fig. 4F**).

Based on these results, we conducted a mutation analysis of MHC-II and the 11-72 mAb at their interaction interface. MHC-II mutants demonstrated equal or enhanced presentation of FR^+^ HEL peptides (**fig. S5B**). Conversely, binding of the 11-72 mAb to Y60A/β and Y65A/β mutants was significantly reduced (**fig. S5C**). The aromatic ring of tyrosine proved crucial for antibody recognition, as substitution with phenylalanine restored antibody binding. A similar pattern was observed with antibody mutants at Y34/H, Y106/H, and Y91/L (**fig. S5D**). It was concluded that not all interaction sites necessarily impact antibody binding. Potential explanations include weak binding or compensatory interactions at other sites.

### iTabs suppress the development of autoimmunity

Experimental autoimmune encephalomyelitis (EAE) serves as an autoimmune disease model for multiple sclerosis, mediated by encephalitogenic CD4^+^ T cells. EAE is induced in SJL/J mice by immunization with the autoantigen, proteolipid protein (PLP) peptide (*20*). It has been reported that the FR^−^ PLP peptide (PLP_139-151_) induces chronic relapsing-remitting EAE more than the PLP peptide with an N-terminal FR (PLP_136-150_)(*21*). We analyzed the Abs in serum from mice immunized with the FR^−^ PLP peptide (PLP_139-151_) or the FR^+^ PLP peptide (PLP_136- 151_) to determine whether iTab was produced. Similar to HEL or SM-P40 peptide, anti-PLP iTab was induced in mice immunized with FR^+^ PLP peptide but not FR^−^ PLP peptide (**Fig. 5A**). Furthermore, immunization with a mutated FR^+^ PLP peptide, in which the pathogenic TCR epitope (PLP_H147K_) is mutated also produced anti-PLP iTab (**Fig. 5A**) (*22*). Similar blocking was observed with monoclonal iTab (clone 2036) obtained from FR^+^ PLP peptide immunized mice (**fig. S6A and S6B**). Activation of five GFP-reporter cells expressing pathogenic anti-PLP TCRs (*20*) was blocked by the monoclonal iTab except for SPL1.1 TCR (**Fig. 5B**).

**Fig. 1.**
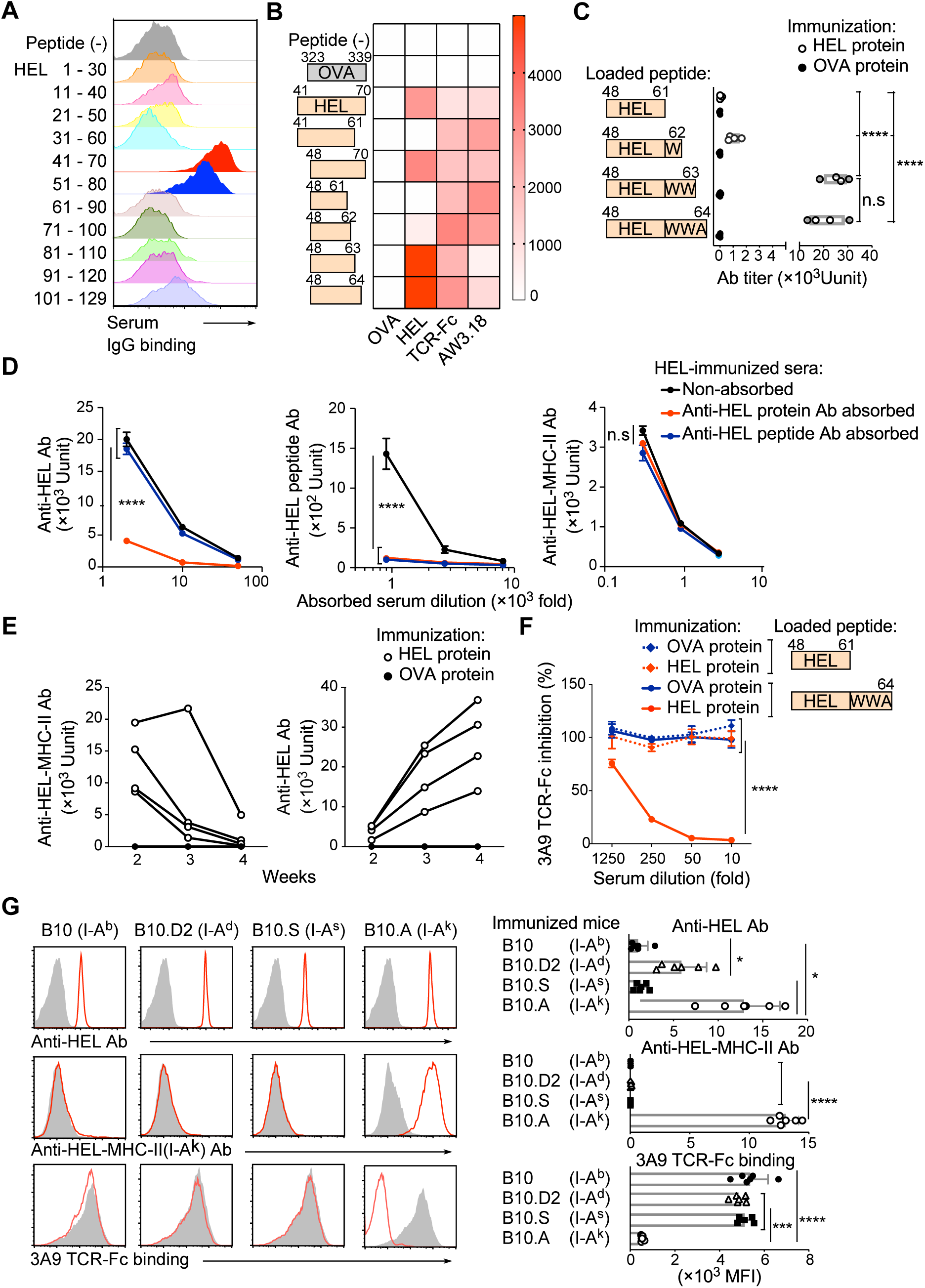
Production of iTabs during immune response to HEL protein. **(A,B)** The binding of serum antibodies (Abs) from HEL protein-immunized mice to HEL peptide-pulsed MHC-II expressing 293T cells was analyzed by flow cytometry (A). Mean fluorescence intensities of Ab binding are presented as a heatmap (B). Serum Abs from OVA-immunized mice, 3A9 TCR-Fc, and Aw3.18 Ab were used as controls. (**C)** Serum Ab titers against HEL peptide-pulsed cells are shown. (n = 4 per group) **(D)** Antibody titers against HEL protein (left), HEL peptide (center), and HEL48-64 peptide-loaded MHC-II expressing cells (right). (n = 3 technical replicates) **(E)** Time course of anti-HEL iTab (left) or anti-HEL Ab (right) titers after HEL protein immunization. Serum from OVA-immunized mice was analyzed as a control. (n = 4 per group) **(F)** Blocking of 3A9 TCR-Fc binding to HEL peptide-loaded MHC-II expressing cells by serum Abs from HEL or OVA protein-immunized mice. (n = 3 technical replicates) **(G)** iTab production in B10 congenic mice. Anti-HEL protein Abs (red line, upper row), anti-HEL iTabs (red line, middle row), and blocking of 3A9 TCR-Fc binding by iTabs (red line, lower row) are shown. (n = 5 per group) Non-immunized mouse sera were used as controls (shaded histogram).Data are presented as the means ± SD [(C), (D), (F) and (G)] Statistical significance was calculated by one-way ANOVA [(C) and (G)] and two-way ANOVA [(D) and (F)] followed by Tukey’s correction; \**p* ≤ 0.05, \*\**p* ≤ 0.01, \*\*\**p* ≤ 0.001, \*\*\*\**p* ≤ 0.0001. All data are representative of three or more experiments.

**Fig. 2.**
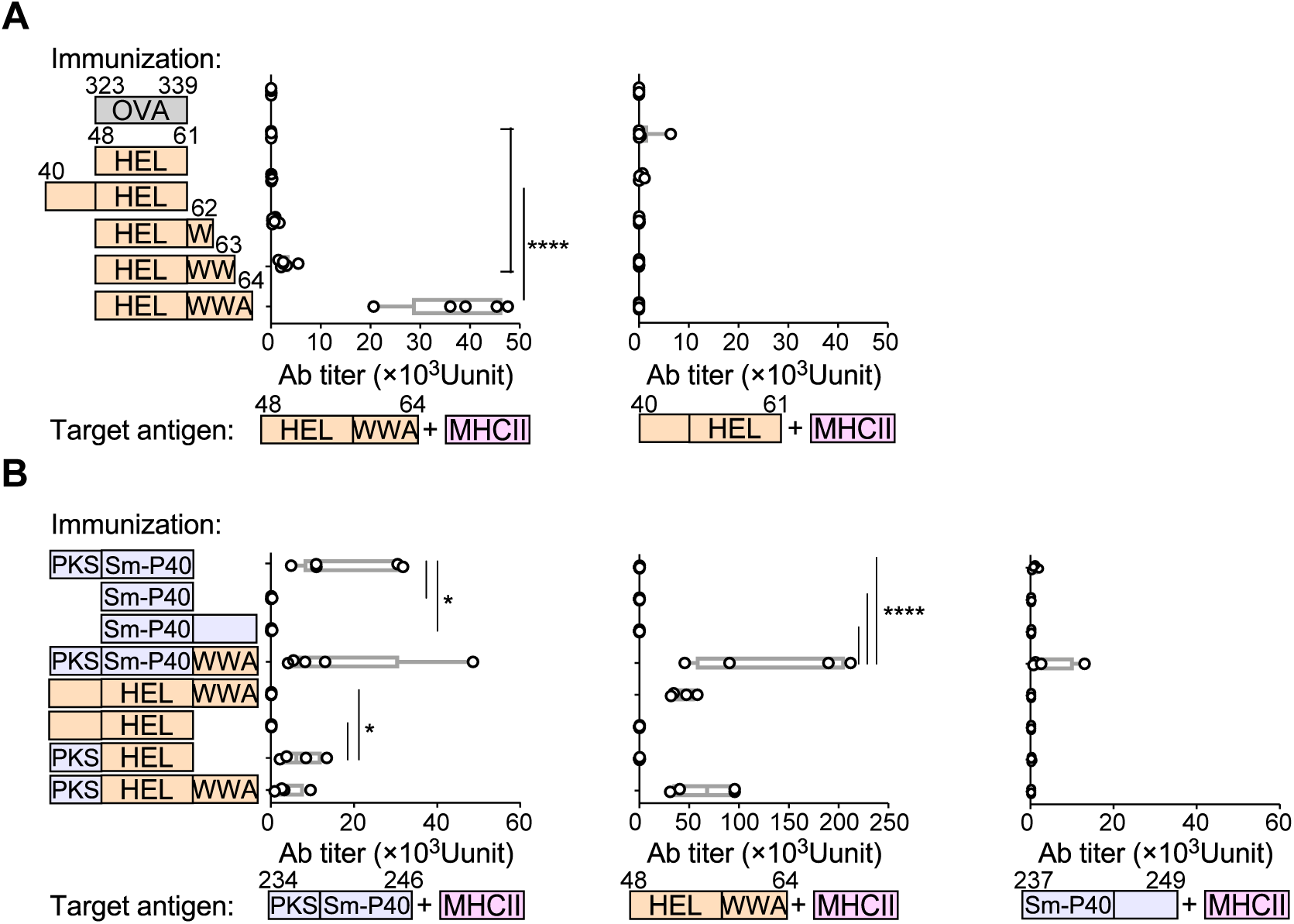
Production of iTabs during immune response to antigen peptides. **(A)** Serum iTab titers induced by immunization with FR^+^ or FR^−^ HEL peptides. Ab titers at different lengths of peptides presented on MHC-II. (*n* = 4-6 mice per group) **(B)** Serum iTab titers induced by immunization with HEL, Sm-P40 or hybrid peptides. iTab titers against FR^+^ Sm-P40 (left), FR^+^ HEL (middle) and FR^−^ Sm-P40 peptide (right)-pulsed MHC-II expressing cells (*n* = 4-5 mice per group). *p* values were determined by one-way ANOVA (A) and (B) with Tukey’s correction was used. Bars represent means ± SD; \**p* < 0.05; \*\**p* < 0.01; \*\*\**p* < 0.001; \*\*\*\**p* < 0.0001. Data represent at least three independent experiments.

**Fig. 3.**
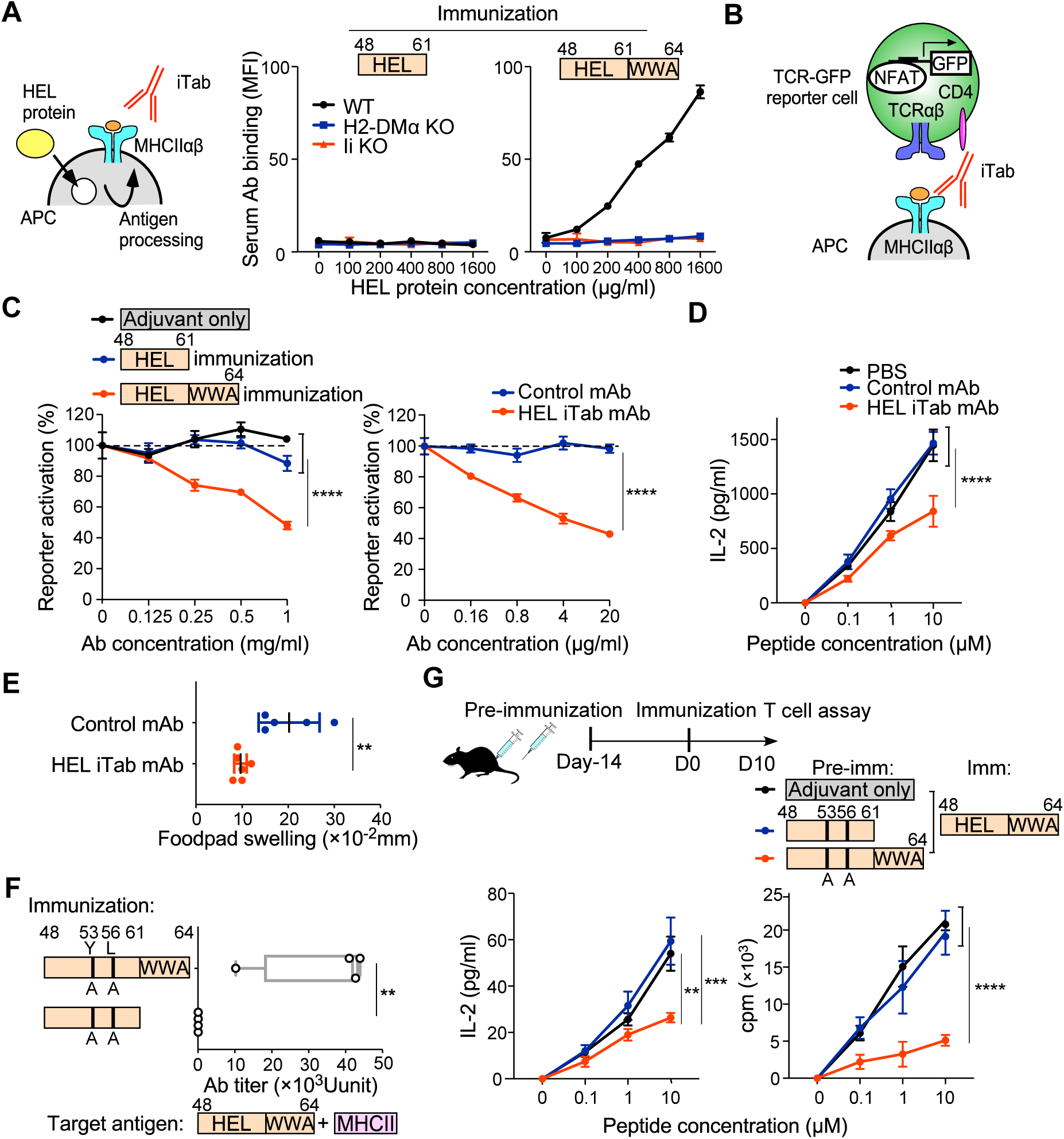
Blocking of antigen-specific T cell response by iTabs. **(A)** Binding of iTabs induced by FR^−^ (left) or FR^+^ (right) HEL peptide immunization was analyzed against WT (black line), H2-DMα (blue line) or invariant chain (red line) knockout cells pulsed with HEL protein (*n* = 3 technical replicates) **(B)** TCR NFAT-GFP reporter system. **(C)** Inhibition of 3A9 TCR reporter cell activation by iTabs from FR^−^ or FR^+^ HEL peptide immunized mice (left) and monoclonal anti-HEL iTab 11-72 (right). (*n* = 3 technical replicates) **(D)** Inhibition of IL-2 production by HEL-specific T cells from mice immunized with FR^+^ HEL peptide by anti-HEL iTab. (*n* = 3 technical replicates) **(E)** Inhibition of DTH response by anti- HEL iTab. (*n* = 5-6 mice per group) **(F)** Production of iTabs by mutated FR^−^ or FR^+^ HEL peptide. (*n* = 4 mice per group) **(G)** Inhibition of HEL-specific T cell response by immunization with mutated FR^+^ HEL peptide. Mice immunized with mutated FR^−^ or FR^+^ HEL peptide were further immunized with wild-type HEL peptides. IL-2 production (left) and T cell proliferation (right) upon FR^+^ HEL peptide stimulation were shown. (*n* = 3 technical replicates) *p* values were determined by two-way ANOVA (C) with Bonferroni’s correction, and two-way ANOVA (D) and (G) with Tukey’s correction, except (E) and (F) where a Student’s *t*-test was used. Bars represent means ± SD; \*\**p* < 0.01; \*\*\**p* < 0.001; \*\*\*\**p* < 0.0001. Data represent at least three independent experiments.

**Fig. 4.**
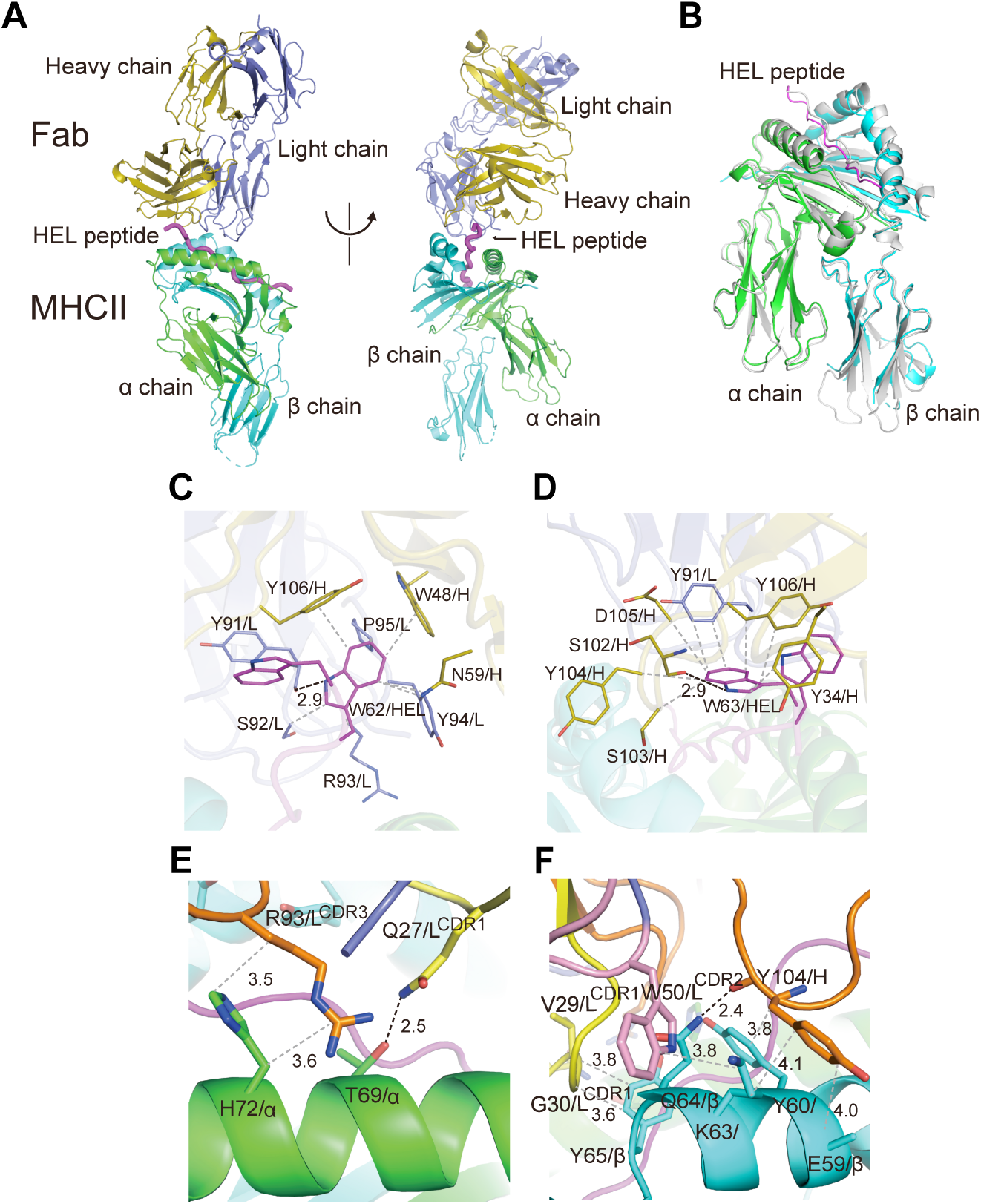
Peptide including FR and MHC-II binding groove as targes of iTab **(A)** Overall structure of the MHC-II-HEL peptide-iTab Fab ternary complex. **(B)** Comparison of the MHC-II structures. Superimposition of the MHC-II-HEL peptide-iTab Fab ternary complex and the murine MHC class II I-A^k^ in complex with HEL peptide (PDB code 1lAK, grey). Fab fragment is omitted here for clarify. The α and β chains of MHC-II, heavy and light chains of Fab fragment, and HEL peptide are colored green, cyan, slate blue, olive, and magenta, respectively. **(C,D)** The interaction of W62 (C) and W63 (D) residues of the HEL peptide and Fab fragment. **(E, F)** The interaction of the α (E) and β (F) chains of MHC-II and Fab fragment. Residues involved in the interaction are shown as stick models. The α and β chains of MHC-II, heavy and light chains of Fab fragment, and HEL peptide are colored green, cyan, slate blue, yellow, and magenta, respectively.CDR1, CDR2 and CDR3 of Fab fragments are colored yellow, pink, and orange, respectively.

**Fig. 5.**
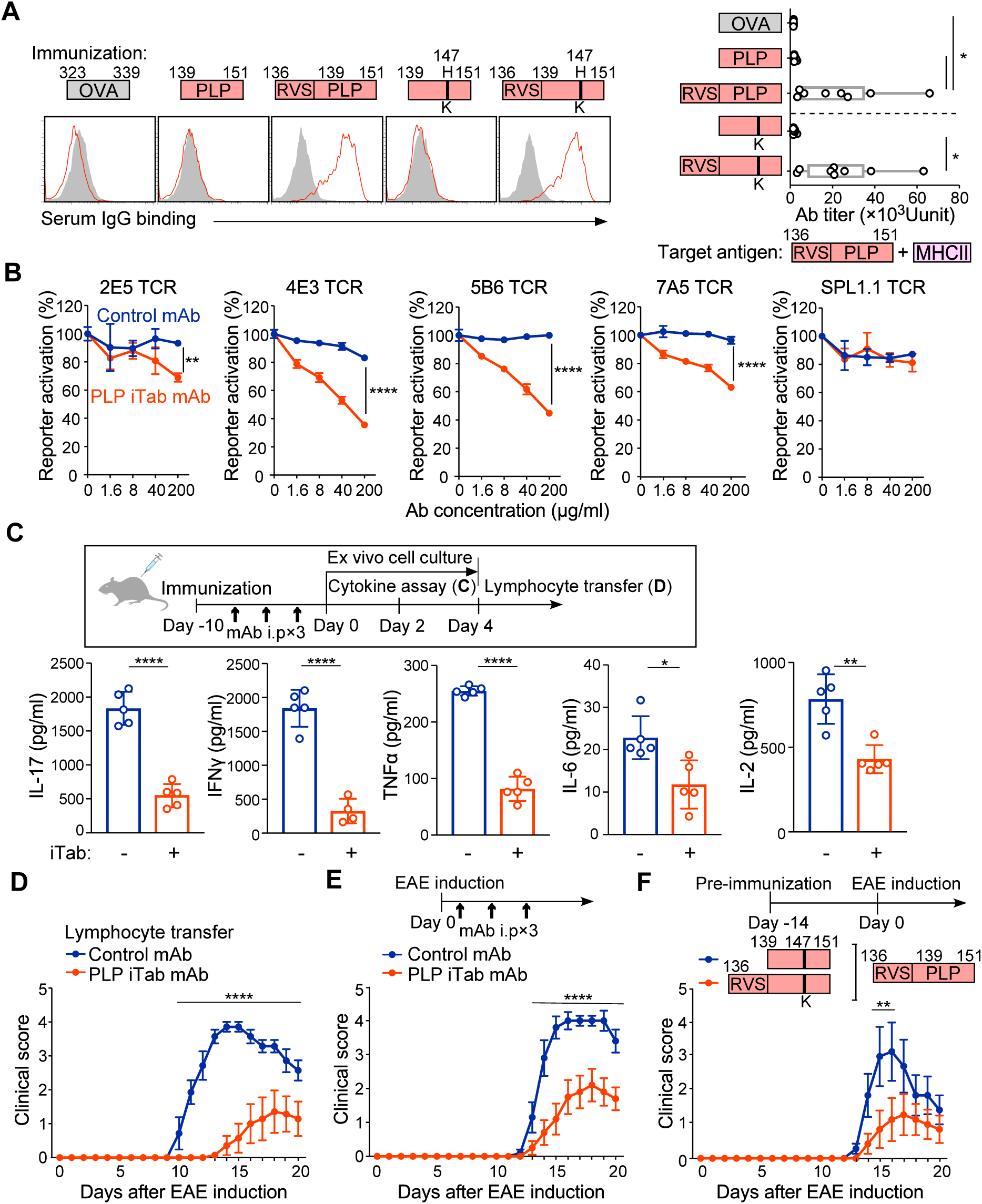
iTabs induced by FR^+^ self-peptide ameliorate experimental autoimmune encephalomyelitis. **(A)** iTab production by immunization with WT PLP, mutated PLP or OVA peptide. PLP- peptide pulsed cells were stained with immunized sera (red line). Control staining was shown as shaded histogram (left). Serum Ab titers against PLP peptide-pulsed cells are shown (right). (*n* =9 mice per group) **(B)** Monoclonal anti-PLP iTab, 2036, blocks the activation of NFAT- GFP reporter cells with PLP-specific TCRs in a dose dependent manner. (*n* = 3 technical replicates) **(C)** iTab treated mice show decreased T cell response to FR^+^ PLP peptide immunization. Inflammatory cytokine production by T cells upon stimulation with PLP peptide was shown (*n* = 5 technical replicates) **(D)** Clinical score of adoptive transfer EAE of ex vivo stimulated lymphocytes. (*n* = 7 mice per group) **(E)** Anti-PLP iTab ameliorates PLP peptide- induced EAE. (*n* = 10 mice per group) **(F)** Anti-PLP iTab-inducing peptide immunization ameliorates EAE. (*n* = 7 mice per group) Data are presented as the means ± SD [(A) to (C)] and ± SE [(D) to (F)]. Statistical significance was tested by one-way ANOVA with Tukey’s correction[(A) to (B)], and two-way ANOVA [(D), (E) and (F)] with Bonferroni’s correction, except for (C) where a Student’s *t* test was used. * *p* ≤ 0.05, \*\**p* ≤ 0.01, and **** *p* ≤ 0.0001. All data are representative of three independent experiments.

To investigate the effect of anti-PLP iTabs on the development of EAE, we analyzed cytokine production in the culture supernatant of lymphocytes from FR^+^ PLP peptide-immunized mice treated with anti-PLP iTabs. Lymphocytes co-cultured with the FR^+^ PLP peptide from anti-PLP iTab-treated mice showed significantly reduced inflammatory cytokine production compared with lymphocytes from mice treated with the isotype control Ab-treated mice (**Fig. 5C**). When T cells from the anti-PLP iTab-treated mice were adoptively transferred to naïve SJL/J mice, T cells from the iTab-treated mice induced significantly milder EAE with delayed onset than T cells from the control mAb-treated mice (**Fig. 5D**). When EAE-induced mice were treated with anti-PLP iTab, the iTab significantly suppressed the progression of EAE (**Fig. 5E**). These results suggested that self-antigen specific-iTabs are involved in the regulation of autoimmunity. Finally, we examined whether immunization with iTab-inducing peptide could prevent EAE.

Mice immunized with the FR^+^ PLP_H147K_ peptide, which induces iTabs without stimulating pathogenic T cells (**fig. S6C**), showed significantly milder EAE symptoms compared to FR^−^ PLP_H147K_ peptide immunized mice in which iTabs are not induced (**Fig. 5F**). These results suggest that iTabs play a role in suppressing self-antigen specific immune response

## Discussion

Although antibodies against the peptide-MHC-II complex can be produced by immunization with a purified peptide-MHC-II complex, it had not been known that iTabs are produced in a general immune response. In this study, we discovered that iTabs are produced during the immune response to several antigens. Because iTabs suppress CD4^+^ T cell response by blocking TCR recognition of the peptide-MHC-II complex, iTabs are involved in the convergence of the immune response or the prevention of excessive immune response.

FR at the N-terminus or C-terminus of antigen peptides was required for iTab production. In general, minimum peptides without FR have been used to analyze CD4 T cells response. Therefore, iTabs have not been observed in most peptide immunization. However, mass spectrometry analyses have demonstrated that many naturally processed peptides presented on MHC-II contain FR (*12, 13*). Although the specific amino acid residues involved in the production of iTab have not been identified, it is important to note that the immunogenicity of peptides varies greatly in the presence or absence of iTabs-inducing FR. Antigen-specific antibody titers generally increase after immunization. However, iTabs titers peaked two weeks after immunization and declined thereafter. This suggests that the mechanism of iTab production is different from the general antibody response to foreign antigens and that iTabs play a role primarily in downregulating the peak immune response.

Structural analysis confirmed that iTabs recognize both FR and MHC-II. Analyses of iTab or MHC-II mutants showed that iTabs recognize both FR and MHC-II. Importantly, peptides lacking FR did not induce iTabs, highlighting the critical role of C-terminal or N-terminal FRs in antigenicity for iTab induction. The binding sites of iTabs differed from those of previously reported TCR-like antibodies that recognize core peptide epitopes presented on MHC-II (*3, 6*). These previously reported TCR-like antibodies were generated by direct immunization with peptide-MHC-II complex, suggesting that these antibodies appear to be generated in a different manner than the iTabs. Furthermore, iTabs blocked most EAE-inducing pathogenic TCR recognition. Optimization of iTab-inducing antigens may lead to the development of more effective methods of specifically suppressing pathogenic autoreactive T cell activation. Our data demonstrate, for the first time, that the induction of iTabs with FR^+^ antigens is a effective strategy to specifically control excessive immune responses observed in autoimmune diseases and allergies.

## Acknowledgements

We are grateful to Nana Yoshikawa, Kaori Oshimo, and Asa Tada for their technical assistance and to Yumi Inaba for her administrative assistance.

## Author Contributions

K.K. performed most of the experiments, analyzed and discussed the data. W.N. discussed the data and assisted with experiments. Ke.K. and K.Y. screened frozen-hydrated samples on the EM grids and collected cryo-EM data. Ke.K. performed EM data processing, structure analysis, model building and structure interpretation. H.T. and S.Y. performed protein sample preparation for cryo-EM, and structural data analysis and discussion. H.A. designed the study. All authors contributed to the writing of the manuscript.

## Author Information

The authors declare no competing financial interests. Correspondence and requests for materials should be addressed to H.A.

## Methods

### Mice

B10.A, B10.D2, B10.S and C57BL/10 congenic mice were purchased from Japan SLC. SJL/J mice were purchased from Charles River Laboratories Japan. All experiments were conducted in accordance with the guidelines of the Animal Research Committee of the Research Institute for Microbial Diseases, Osaka University.

### Cells

The human embryonic kidney 293T cell and LK35.2 cell expressing I-A^k^ were obtained from the RIKEN Cell Bank and the American Type Culture Collection. The invariant chain (Ii) or H-2M alpha deleted LK35.2 cells were generated by CRISPR-Cas9 system using pX330 vector (Addgene: ID42230) inserted into the primers (Ii:5’-caccgtacaccggtgtctctgtcc-3’ and 5’- aaacggacagagacaccggtgtac-3’, H-2M alpha 5’-caccgattcccaacatagggctct -3’ and 5’-aaacagagccctatgttgggaatc-3’). Each cell line was tested regularly for *Mycoplasma* contamination using PCR.

### Plasmids and Transfection

Plasmids for MHC class II molecules were constructed as previously described (*10*). The cDNAs were cloned into the pME18S expression vector. 293T cells were transiently co-transfected with MHC class II molecules and pMxs-GFP using PEI Max (Polysciences) and GFP-positive cells were analyzed 3 days post-transfection. The plasmids for the 3A9 single chain TCR-Fc fusion protein were generated by cDNA from the 3A9 T cell hybridoma with 3×(GGGS)linkers between the extracellular domains of the α and β chains. The construct was cloned into the pME18S expression vector which contains the human IgG1 constant region. Plasmids for mouse CD4 (accession No.: NM_013488), and TCR α and β genes of 3A9, 2E5, 4E3, 5B6, 7A5 and SPL1.1 were synthesized (Integrated Device Technology) according to the published TCR gene sequence and cloned into pMxs retroviral vector (*20*). The generation of these CD4 and TCRαβ stable transfectants was achieved with PLAT-E retroviral packaging cells with an amphotropic envelope.

### Peptides

The HEL peptide library, which overlapped by 10 residues was purchased from SCRUM Inc. Other peptides were obtained from GeneScript, with a purity of > 90% as determined by high-performance liquid chromatography. The epitopes encoded in the different monomeric constructs used here include; OVA_323-339_(ISQAVHAAHAEINEAGR), HEL_41- 70_(QATNRNTDGSTDYGILQINSRWWCNDGRTP), HEL_41-61_(QATNRNTDGSTDYGILQINSR), HEL_48-70_(DGSTDYGILQINSRWWCNDGRTP), HEL_48-61_(DGSTDYGILQINSR), HEL_48-62_(DGSTDYGILQINSRW), HEL_48-63_(DGSTDYGILQINSRWW), HEL_48-64_(DGSTDYGILQINSRWWA), biotinylated HEL_48-64_(BioGSGSDGSTDYGILQINSRWWA), Sm-P40_234-246_(PKSDNQIKAVPAS), Sm-P40_237-246_(DNQIKAVPAS), Sm-P40_237-249_(DNQIKAVPASQAL), Sm-P40_234-246-WWA_(PKSDNQIKAVPASWWA), _PKS-_HEL_52-61_(PKSDYGILQINSR), _PKS-_HEL_52-64_(PKSDYGILQINSRWWA), HEL_48-61(53,56A)_ (DGSTDAGIAQINSR), HEL_48-64(53,56A)_ (DGSTDAGIAQINSRWWA), PLP_136-151_(RVSHSLGKWLGHPDKF), biotinylated PLP_136- 151_(BioGSGSRVSHSLGKWLGHPDKF), PLP_139-151_(HSLGKWLGHPDKF), PLP_136-151(H147K)_ (RVSHSLGKWLGKPDKF), PLP_139-151(H147K)_ (HSLGKWLGKPDKF)._We employed the HEL_48-64_ peptide, in which the C-terminal cysteine was substituted with alanine, in order to inhibit the formation of S-S bonds in this paper.

### Immunization

Mice were immunized subcutaneously (s.c.) with 100μg of HEL (L6876, Sigma-Aldrich), chicken ovalbumin (A5503, Sigma-Aldrich), or 30nmol of peptides in Complete Freund’s adjuvant (F5881, Sigma-Aldrich). Pre-immunization of mutated peptides was conducted using Incomplete Freund’s adjuvant (263910, BD Difco). Serum samples were collected from immunized six to eight-week-old female mice after a period of 2-4 weeks. B cell hybridomas were established according to standard protocols. Briefly, lymphocytes from lymph nodes were fused to P3U1 cells to generate monoclonal antibodies, which were then screened using MHC- II-transfectants pulsed with peptide by flow cytometry. Antibodies from serum or monoclonal antibodies were purified using Protein A and G sepharose (GE Healthcare) on a Profinia protein purification system (Bio-Rad) and gel filtration on an AKTA pure 25 (GE Healthcare)

### Flow cytometry

Antigen-presenting cells (APCs) were prepared by stimulating them with peptides or proteins for either overnight or 36 hours. The cells were stained with immunized mouse serum at a dilution of 1:100-1000 and then reacted with allophycocyanin (APC)-conjugated anti-mouse IgG (715-136-151, Jackson ImmunoResearch Laboratories) or Alexa Fluor 647-conjugated anti-mouse IgG1(A21240, Invitrogen) for LK35.2 cell staining as a secondary antibody. The Aw3.18 antibody was purified from the supernatant of a hybridoma culture (CRL-2826, ATCC). Antibody titers were determined by serial dilution of positive serum. Anti-HEL or HEL peptide antibodies in serum were removed by Streptavidin (SA) conjugated sepharose beads (GE Healthcare) fused with biotinylated antigen. For the detection of anti-peptide antibodies or anti-HEL antibodies, SA (Z02043, Genscript) was labelled with aldehyde/ sulfate latex beads (A37304, Invitrogen), and then biotinylated peptide or protein was mixed. The binding of the human receptor-Fc fusion 3A9 TCR-Fc was examined by premixed with an APC-conjugated anti-human IgG Fc antibody (109-136-098, Jackson ImmunoResearch Laboratories). The TCR-Fc competitive assay was conducted by reacting the diluted serum before the premixed-TCR-Fc, followed by staining. APC-conjugated anti-mouse CD4 (17-0041-83, eBioscience) was employed in the reporter assay. For intracellular staining, cells were fixed and permeabilized using the BD Cytofix/Cytoperm Fixation/Permeabilization Solution Kit (554714), then stained with anti-mouse H2-DM (552405, BD Biosciences), Ii (151002, Biolegend) and APC-conjugated anti-rat IgG (712-136-153, Jackson ImmunoResearch Laboratories). Flow cytometric analyses were conducted using either a FACSCalibur or a FACSVerse flow cytometer (BD Biosciences). The data were subsequently analyzed using FlowJo v10 software (FlowJo, LLC).

### Reporter Assay

The TCR reporter cells were mouse T cell hybridomas that had been stably transfected with NFAT-GFP and FLAG-tagged DAP12, as previously described (*23, 24*). The original hybridoma TCR αβ chains were depleted using the CRISPR-Cas9 system with the pX330 vector inserted into the primers (α chain; 5’-CACCGTGCCGAAAACCATGGAATC-3’ and 5’-AAACGATTCCATGGTTTTCGGCAC-3’, βchain; 5’- CACCGAGAAATGTGACTCCACCCA-3’ and 5’-AAAC TGGGTGGAGTCACATTTCTC-3’). Following cell depletion, mouse CD4 was stably transfected, and each TCR was introduced into the CD4-positive cells. The cells were sorted using a Cell Sorter (SH800Z, Sony). The TCR reporter cells (96-well plate at 2 × 10^4^ cells per well) were co-cultured with HEL protein-pulsed LK35.2 cells or PLP_136-151_ peptide-loaded 293T cells transfected with I-A^s^ (4 × 10^4^ cells per well) and supplemented with or without antibodies for 16 hours. The expression of GFP was analyzed using flow cytometry.

### In vitro Proliferation and IL-2 secretion assays

CD4^+^ T cells (2 × 10^5^ cells per well) were isolated from peripheral lymph nodes of peptide - immunized mice using BioMag anti-mouse IgG (QIAGEN) and a mouse CD4 T cell isolation kit (Miltenyi Biotec). Splenocytes as APC (6 × 10^5^ cells per well) from wild-type mice were treated with an Ack buffer and subsequently treated with mitomycin C. The cells were incubated with HEL_48-64_ peptides for 72 hours in Advanced RPMI 1640 (Invitrogen) supplemented with 1% FBS (Hyclone), Glutamax (Invitrogen) and penicillin/streptomycin (Nacalai tesque), and pulsed with 1 μCi (^3^H) thymidine for 18 hours before harvesting and reading on a beta counter reader (Perkin Elmer). Supernatants were collected 24 hours later for the measurement of IL-2 by ELISA ready-set-go reagent set (eBioscience).

### Delayed-type hypersensitivity (DTH) assays

Ten days post-immunization of HEL_48-64_ peptide, mice were challenged with 30 nmol of the same peptide, which was injected intradermally into the footpad. The right footpad was injected with the peptide, while the left footpad was injected with vehicle (PBS) as a baseline control. The degree of swelling was quantified 24 hours post-challenge using a Thickness Gauge (SM- 112, TECLOCK). Monoclonal Abs (1 mg per injection) were administered intraperitoneally (i.p.). on day 3 and 6.

### Sample preparation for cryoEM analysis

The construction of soluble MHC class II molecules has been previously described. The I-A^k^ alpha plasmids were fused with a GSSGSSG-linker, TEV protease cleavage site, leucine zipper and His-tag at the C-terminus, and HEL_48-64_ peptide was bound to the I-A^k^ beta plasmids with a GGGGSLVPRGSGGGGSGS-linker, which was attached via a GSSGSSG-linker, TEV protease cleavage site, leucine zipper and a Flag-tag at the C-terminus. The plasmids were subsequently inserted into the pcDNA3.4 expression vector. The recombinant protein was obtained via the Expi293 expression system (Thermo Fisher Scientific). Monoclonal Abs were obtained from hybridoma cells by digestion to the Fab fragment using immobilized papain (20341, Sigma-Aldrich). The purified HEL peptide-MHCII protein was mixed with the Fab fragment, and incubated on ice for 30 min. The mixture was loaded onto a Superdex 200 10/300 column equilibrated with 20 mM Tris-HCl buffer (pH 8.0) containing 150 mM NaCl and was eluted with the same buffer. Fractions containing the complex were collected and concentrated to approximately 4 mg/ml by ultrafiltration.

### Cryo-EM data collection

Holey carbon film-coated copper grids (Quantifoil R1.2/1.3 Cu 200 mesh; Microtools GmbH, Berlin, Germany) pretreated by Au sputtering were glow-discharged for 10 s using an ion coater (JEC-3000FC; JEOL, Tokyo, Japan). A solution of 4.0 µL (1.0 mg/mL) of the anti-HEL iTab, suspended in a buffer containing 20 mM Tris-HCl (pH 8.0) and 100 mM NaCl, was applied to the grid and blotted with filter paper for 6 s. The grid was then immediately plunged into a bath of cooled ethane using a FEI Vitrobot Mark IV (Thermo Fisher Scientific, Waltham, MA, USA) under 100% humidity at 4 °C. Subsequently, the grids were examined using a CRYO ARM 300 electron microscope (JEOL)(*25*), equipped with a cold-field emission gun and an in-column energy filter with a slit width of 20 eV. Dose-fractionated images were recorded using a K3 Summit direct electron detector (AMETEK, Berwyn, PA, USA) in counting mode, with a dose rate of ∼ 99 e^−^ Å^−2^ across 100 frames. All images were collected using SerialEM(*26*) with AI-assisted hole detection via yoneoLocr(*27*), at a nominal magnification of 100,000×, corresponding to a pixel size of 0.495 Å. The defocus range was estimated to −0.8 µm to −5.0 µm. A total of 17,175 movies were acquired.

### Cryo-EM data processing

Data processing was performed using CryoSPARC 4.2.0(*28*) and Relion 3.1.4(*29*). The collected movies were aligned, dose-weighted, and averaged through motion correction implemented in CryoSPARC. The contrast transfer function (CTF) was estimated using Ctffind-4.1.13(*30*). Subsequently, micrographs with an estimated resolution below 10 Å were excluded from the data processing. A total of 1,327,931 particles in 12,394 micrographs were selected through Blob picker and subjected to the first two-dimensional (2D) classification, template picker, and second 2D classification. Then, an initial three-dimensional (3D) map (C1 symmetry) was reconstructed through ab-initio reconstruction in CryoSPARC. Following homogeneous and un-uniform refinement yielded a 3D map at 3.7 Å resolution. The micrographs and 3D map obtained were exported to RELION 3.1.4, where particles were picked up again using Topaz and 3D reference picking due to a tendency for preferred orientations of the particles. Subsequent 2D and 3D classification in RELION yielded a total of 786,526 particles. Several rounds of 3D refinement, CTF refinement, and Bayesian polishing resulted in a 3D map at 2.91 Å resolution. However, the obtained 3D map was highly disordered, making model building difficult. Therefore, the selected particles and the 3D map were imported back into CryoSPARC, and subjected to 3D classification (20 classes) again, resulting in the selection of 50,032 high-quality particles. Additionally, 3D flexible refinement (3DFlex)(*29*) improved the disorder of the 3D map, resulting in a 3D map at 3.09 Å resolution based on the gold-standard Fourier Shell Correlation cutoff (0.143) criterion, through post processing in RELION. Furthermore, an enhanced map was generated using EMready(*31*), a program for improving the interpretability of the 3D map by leveraging similarity and correlation-guided deep learning, which was used as a reference for model building.

### Atomic model building

An initial model was generated using the structure of murine MHCII class II I-A^k^ with a peptide from hen egg lysozyme (PDB code: 1IAK(*19*)) and H-2 class II histocompatibility antigen (PDB code: 2PXY(*32*)) through the SWISS-MODEL server (https://swissmodel.expasy.org/). These models were fitted into the map using UCSF Chimera(*33*). The fitted initial model was manually adjusted with COOT(*34*) to better fit the 3D map, followed by refinement using Phenix (*35*) and REFMAC5 in CCP-EM(*36, 37*). The refinement statistics of the final model were obtained using the comprehensive validation program in Phenix.

### Data availability

Atomic coordinates and cryo-EM maps for the reported structure of anti-HEL iTab were deposited in the Protein Data Bank under accession code 9L1L, and in the Electron Microscopy Data Bank under accession code EMD-62748.

### Cytokine secretion assay and adoptive transfer of EAE

Female SJL/J mice, aged 10-14 weeks, were immunized s.c. with 100μl of a CFA emulsion containing 200μg of *Mycobacterium tuberculosis* H37Ra (Difco Laboratories) and 15nmol of PLP_136-151_ peptide. Monoclonal anti-PLP iTab (1mg per injection) were administrated i.p. on days 0,4 and 8. Cytokine release was evaluated by culturing lymphocytes from lymph nodes on day 10 in vitro with 0.1 μM of PLP_136-151_ peptide or an irrelevant peptide (OVA_323-339_) for 48 hours. Supernatants were collected and analyzed using a mouse Th1/Th2/Th17 cytokine kit (BD Bioscience) by flow cytometry. lymphocytes were cultured for a further additional 48 hours, after which 1 × 10^7^ cells were transferred to naïve SJL/J mice. The animals were evaluated daily for clinical signs of disease as follows: (0) No disease; (0.5) Loss of tail tonus; (1) Complete tail paralysis; (2) Tail paralysis with hind limb weakness; (3) Complete hind limb paralysis; (4) Hind limbs paralyzed, weakness in forelimbs; (5) Tetraplegia; (6) Death.

### iTab or Peptide therapy in EAE

EAE was induced by immunization with 100μl of a CFA emulsion containing 250μg of *Mycobacterium tuberculosis* H37Ra and 50nmol of PLP_136-151_ peptide. In iTab therapy, Monoclonal anti-PLP iTab (1mg per injection) were administrated i.p. on days 0,4 and 8. Two weeks before the onset of EAE, a pre-immunization procedure was conducted in which 50nmol of PLP peptide, which mutated for the pathogenic TCR recognition site at H147K, was administered s.c. with IFA.

### Statistical analysis

Data were compared by Student’s *t*-test, one-way or two-way ANOVA tests. *p* values < 0.05 were considered statistically significant.

## Extended Data Figure legends

**Fig. S1.**
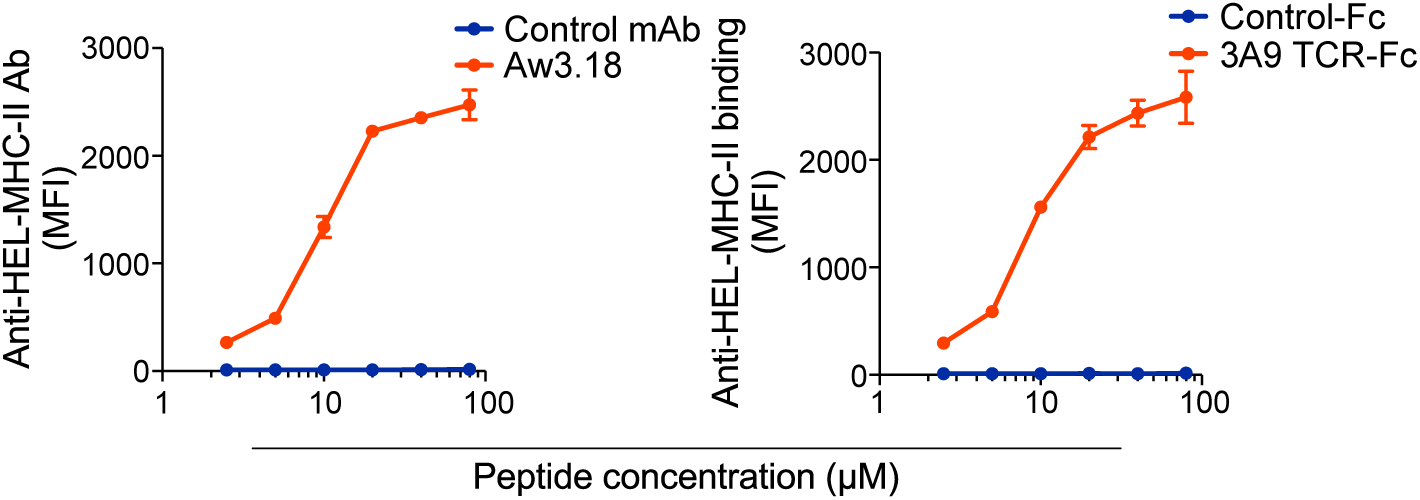
HEL peptide presentation by MHC-II was detected with Aw3.18 and 3A9 TCR-Fc. MHC-II expressing 293T cells were pulsed with different concentrations of HEL_48-61_ peptide and were stained with Aw3.18 Ab (left) or 3A9 TCR-Fc fusion protein (right). Data represent the mean ± SD (*n* = 3 technical replicates). The experiments were replicated twice.

**Fig. S2.**
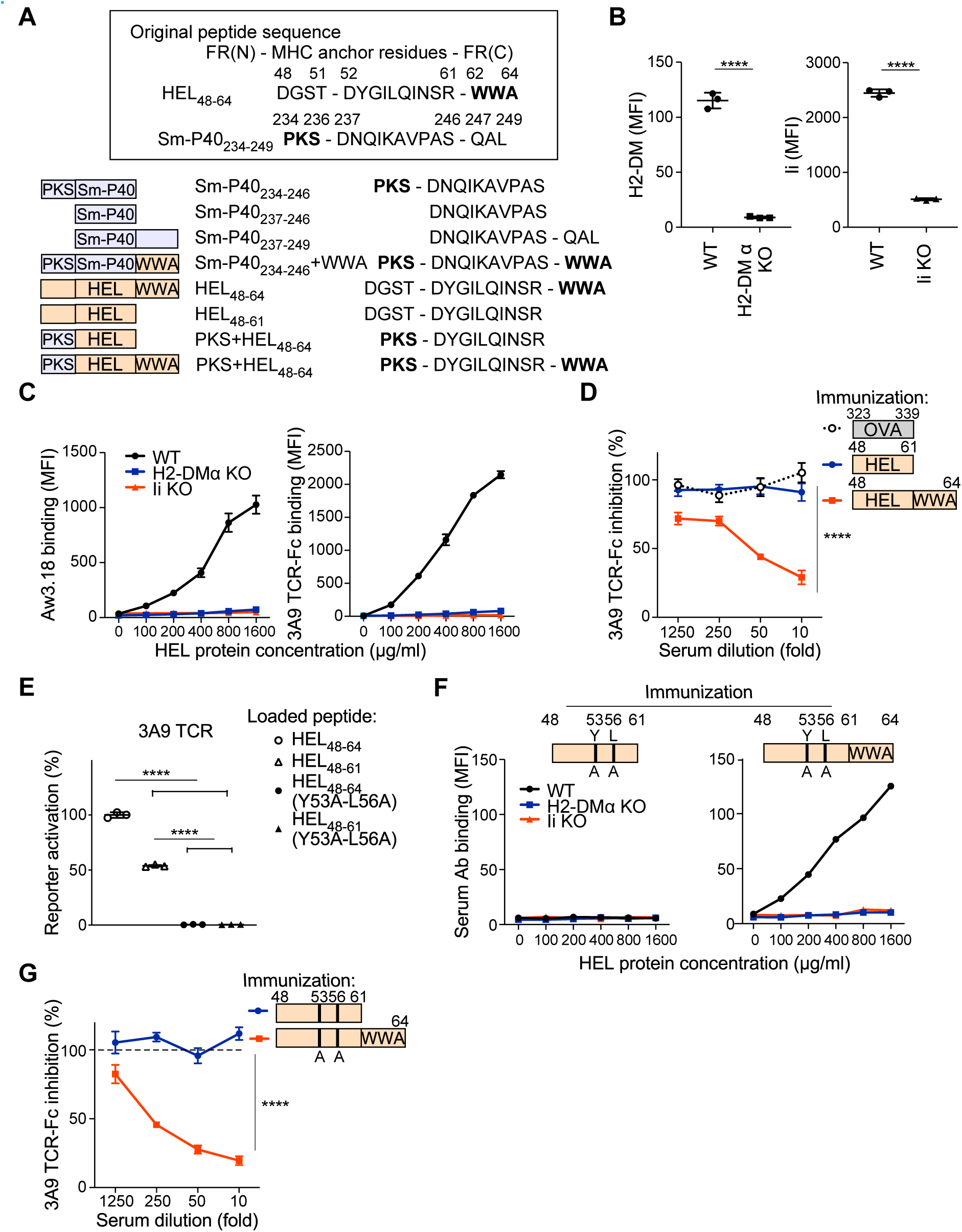
Analysis of Abs in serum immunized with HEL peptide. (A)Peptide sequence of HEL, Sm-P40 and hybrid peptides. iTab inducible FRs were shown in bold. (B) Expression levels of H2-DM or Ii in knockout APCs. (C) Binding of Aw3.18 and 3A9 TCR-Fc against WT LK35.2(black line), H2-DMα (blue line) or invariant chain (red line) knockout cells pulsed with HEL protein. (D) Blocking of 3A9 TCR-Fc binding by Abs induced by immunization with FR^−^ or FR^+^ HEL peptide. (E) Activation of 3A9 TCR-expressing reporter cell with WT and mutated peptide. (F) Binding of serum Abs immunized with mutated FR^−^ or FR^+^ HEL peptides against WT LK35.2(black line), H2-DMα (blue line) or invariant chain (red line) knockout cells pulsed with HEL protein. (G) Blocking of 3A9 TCR-Fc binding by Abs induced by mutated FR^−^ or FR^+^ HEL peptide immunization. *p* values were determined by one-way ANOVA[(E)], two-way ANOVA[(D) and (G)] with Tukey’s correction, except for (B) where a Student’s *t*-test was used; **** *p* ≤ 0.0001. All data represent the mean ±SD (*n* = 3 technically independent samples). The experiments were performed more than twice.

**Fig. S3.**
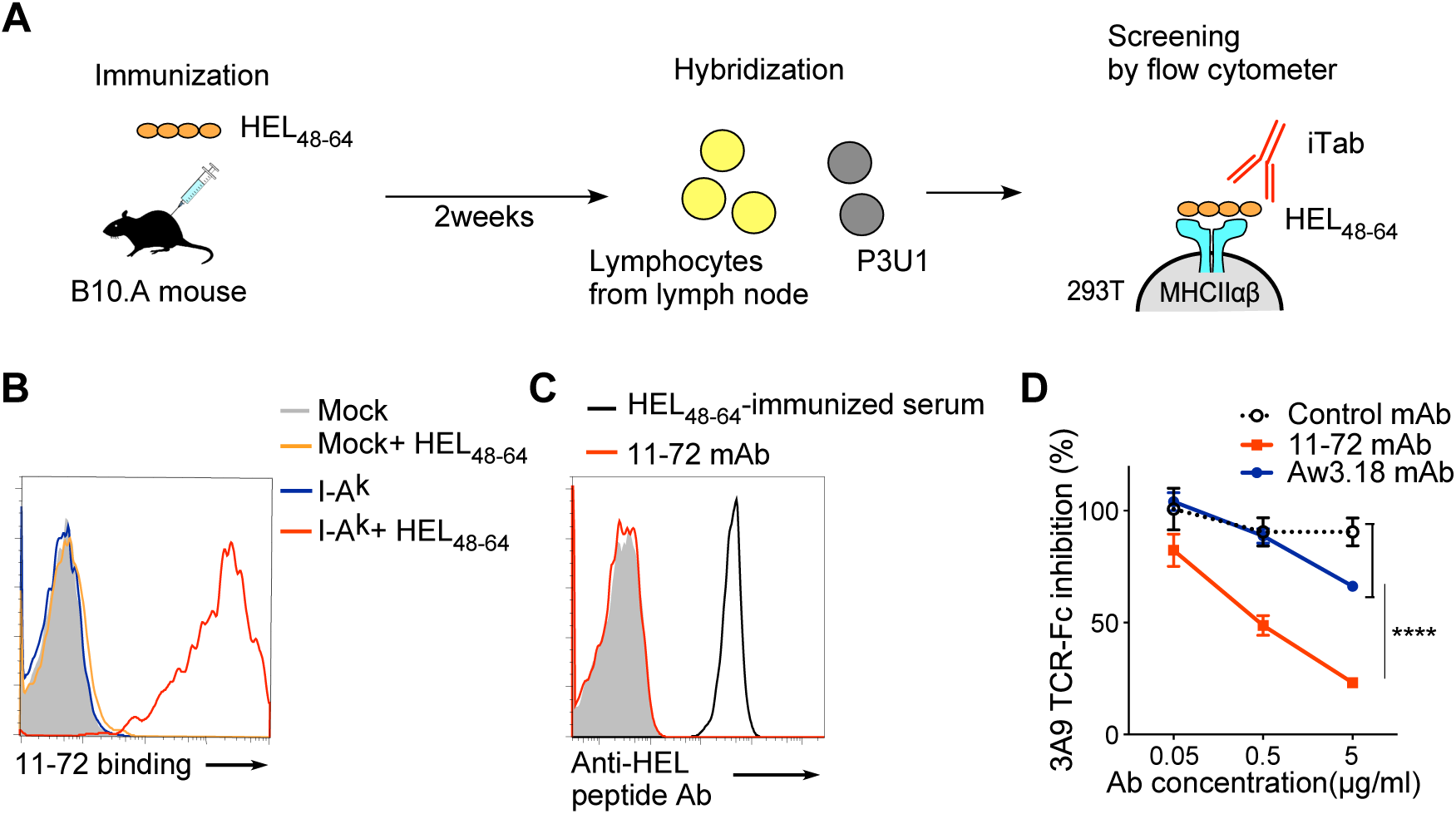
Generation of monoclonal anti-HEL iTab 11-72. Ab concentration(µg/ml) (A) Protocol to generate monoclonal anti-HEL iTabs. (B) Binding of anti-HEL iTab 11-72 to HEL_48-64_ peptide-pulsed cells. (C) Binding of 11-72 iTab to HEL_48-64_ peptide-coated latex beads (right). (D) Blocking of 3A9 TCR-Fc binding by 11-72 or Aw3.18 mAb. *p* values were determined by two-way ANOVA with Tukey’s correction; **** *p* ≤ 0.0001. Data represent the mean ± SD [(D)] (*n* = 3 technically independent samples). The experiments were replicated twice.

**Fig. S4.**
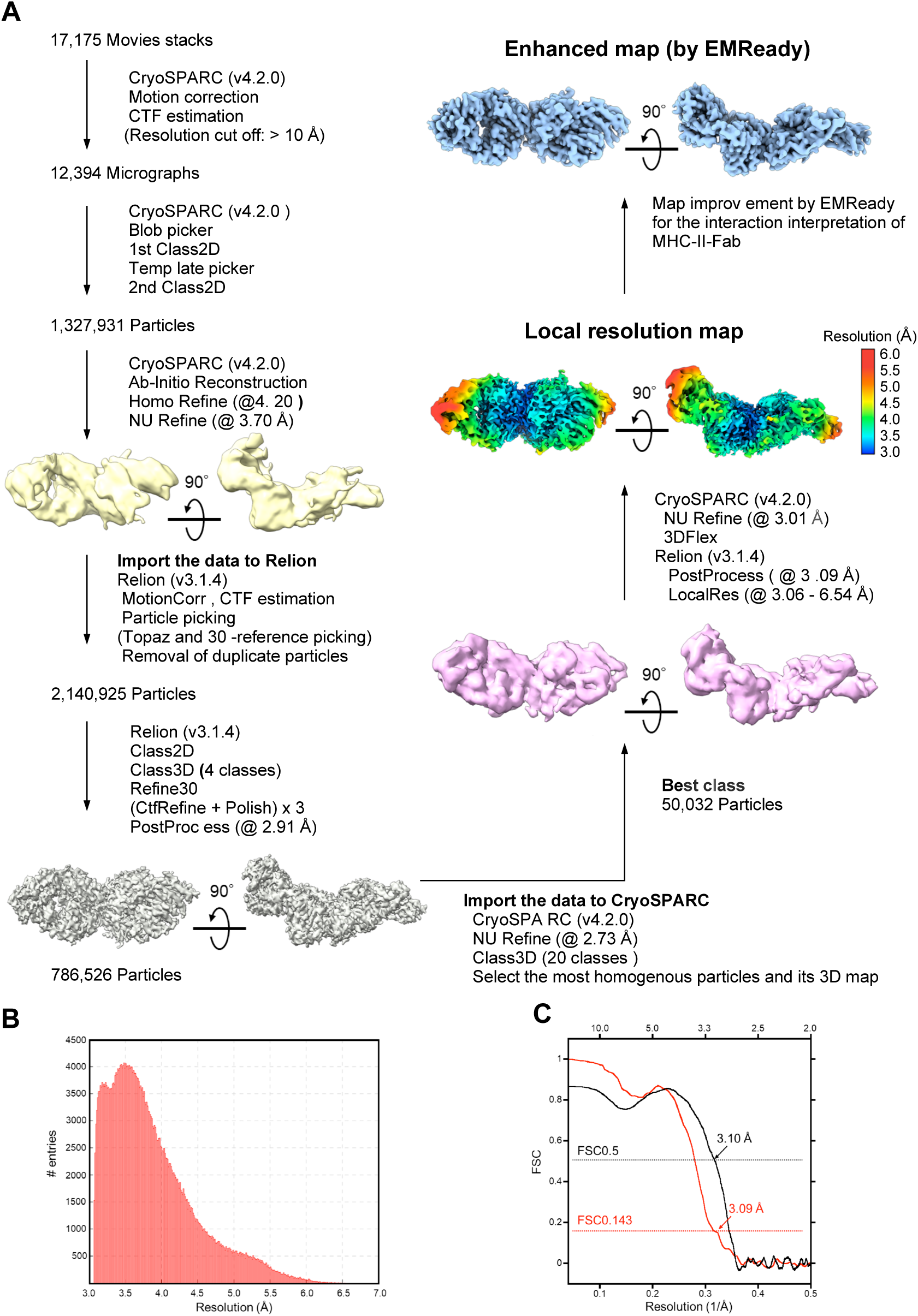
Data collection and image processing workflow of the anti-HEL iTab. (A) Overview of the data processing workflow. The resolution was estimated based on the gold standard Fourier shell correlation (FSC) criteria of 0.143. (B) Histogram of local resolution of the 3D map. (C) FSC curves for 3D reconstruction of the 3D map and the refined model versus the overall 3.1 Å map. Red, gold-standard curve with a value of 0.143 at 3.7 Å resolution; red, FSC curve calculated between the 3D map and the refined structure model of the anti-HEL iTab.

**Fig. S5.**
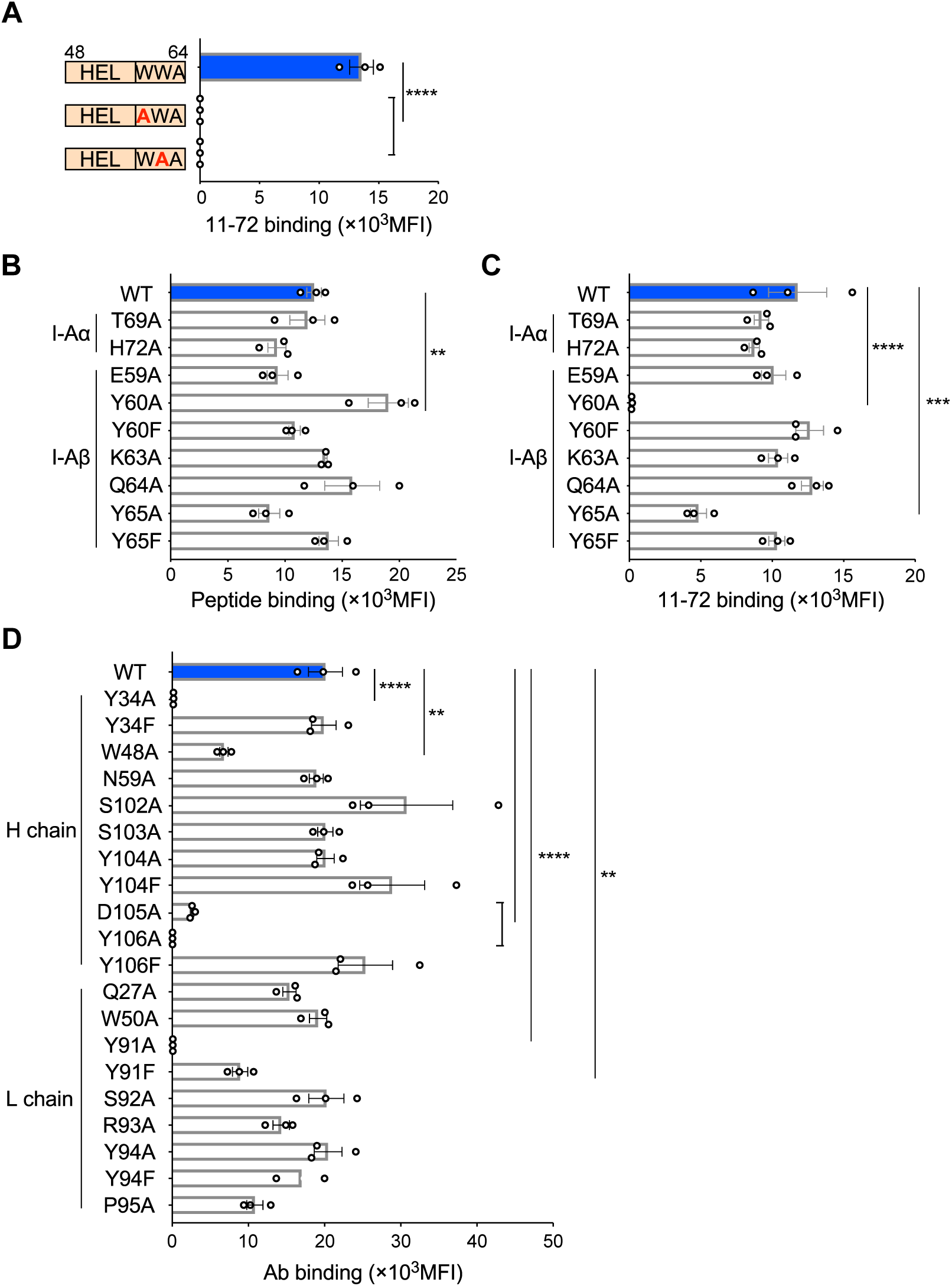
Mutational analysis of 11-72 mAb binding to HEL peptide-pulsed MHC-II. (A) Binding of 11-72 mAb to HEL peptides presented on MHC-II. (B) HEL_48-64_ peptide presentation by MHC-II mutants. (C) 11-72 mAb binding to HEL_48-64_ peptide presented on MHC-II mutants. (D) Binding of 11-72 mAb mutants to HEL_48-64_ peptide pulsed MHC-II. *p* values were determined by one-way ANOVA with Dunnett’s correction; ** *p* ≤ 0.01, *** *p* ≤ 0.001 and **** *p* ≤ 0.0001. Data represent the mean ± SE [(A)-(D)] (*n* = 3 technically independent samples). The experiments were replicated three times.

**Fig. S6.**
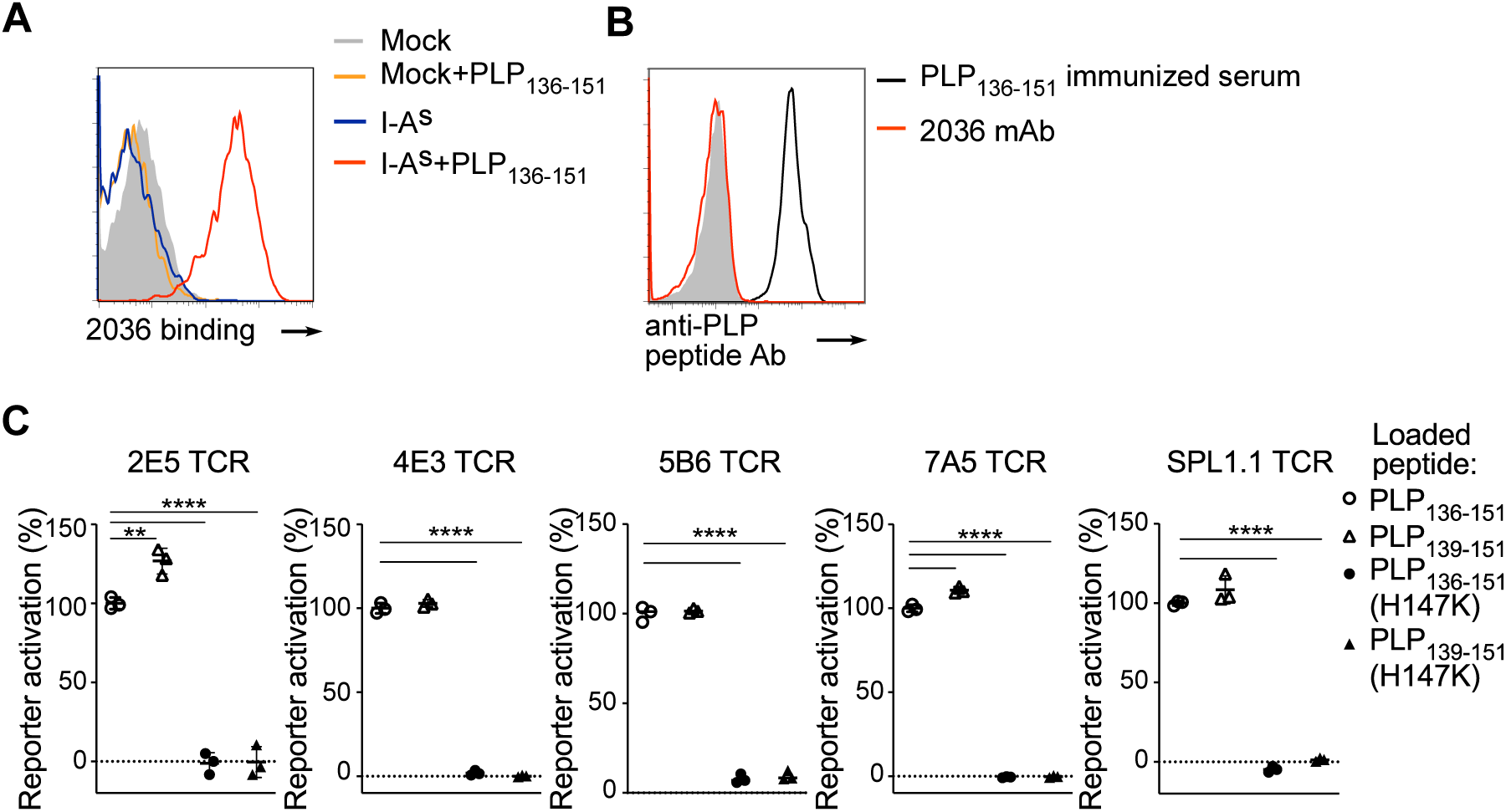
Specificity of monoclonal anti-PLP iTab 2036 and PLP specific-TCR reporter cells. (A) Binding of anti-PLP iTab 2036 to PLP_136-151_ peptide pulsed on MHC-II, I-A^s^, expressing cells. (B) Binding of 2036 iTab to PLP_136-151_ peptide coated beads. (C) Activation of PLP-specific TCR reporter cells to WT and mutant PLP peptides. *p* values were determined by one-way ANOVA with Bonferroni’s correction; ** *p* ≤ 0.01 and **** *p* ≤ 0.0001. Data represent the mean ± SD (*n* = 3 technically independent samples). The experiments were performed more than twice.

**Table S1.** Cryo-EM data collection, refinement, and validation statistics.

|  | anti-HEL AiTab<br>(EMDB- 62748)<br>(PDB 9L1L) |
| --- | --- |
| **Data collection and processing** |  |
| Microscope | CRYOARM 300 |
| Imaging device | K3 summit |
| Nominal magnification | x100,000 |
| Voltage (kV) | 300 |
| Data collection | SerialEM<sup>27</sup>, yoneoLocr<sup>28</sup> |
| Imaging mode | Counting |
| Electron exposure (e<sup>-</sup>/Å<sup>2</sup>) | ~ 99 |
| Frame per movie | 100 |
| Defocus range (μm) | -0.2 – -5.0* |
| Original pixel size (Å) | 0.495 |
| No. of total images sets | 17,175 |
| Pixel size of final map | 0.990 |
| Symmetry imposed | C1 |
| Initial particle images (no.) | 2,659,032 |
| Final particle images (no.) | 50,032 |
| Map resolution (Å) | 3.09 |
| FSC threshold | 0.143 |
| Map sharpening *B* factor (Å<sup>2</sup>) | -53.77 |
| **Refinement** |  |
| Initial model used (PDB code) | 1IAK, 4HC1 |
| Model resolution (Å) | 3.10 |
| FSC threshold | 0.5 |
| Model resolution range (Å) | 247.5 – 3.09 |
| Model composition |  |
| Non-hydrogen atoms | 6,383 |
| Protein residues | 800 |
| Ligands | - |
| *B* factors (Å<sup>2</sup>) |  |
| Protein | 175.1 |
| Ligand | - |
| R.m.s. deviations |  |
| Bond lengths (Å) | 0.007 |
| Bond angles (°) | 1.241 |
| Validation |  |
| MolProbity score | 1.95 |
| Clashscore | 8.80 |
| Poor rotamers (%) | 0.44 |
| Ramachandran plot |  |
| Favored (%) | 92.28 |
| Allowed (%) | 7.07 |
| Disallowed (%) | 0.65 |

\*Defocus range was estimated by CTFFIND4<sup>32</sup>.

